# From amplicons to strains: The limitations of metabarcoding as criterium in the selection process of biocontrol strains against the pome fruit pathogen *Neonectria ditissima*

**DOI:** 10.1101/2023.12.05.570079

**Authors:** Lina Russ, Georgina Elena Jimenez, Jean Luc van den Beld, Els Nijhuis, Jürgen Köhl

## Abstract

European canker, caused by the fungal pathogen *Neonectria ditissima*, causes severe economic losses in apple production and conventional control measures are not sufficiently effective. Recently, parts of the endophyte community have been suggested to play a role in the response of the host to the pathogen, potentially leading to higher resistance of apple cultivars to disease outbreaks. In addition, advances on biologically controlling the disease have been booked by the application of fungi isolated at the boundary of cankered and healthy wood tissue. In this study we sought to evaluate if and how metabarcoding analysis can support decisions on selection of biological control agents in a two-steps process: first we profiled fungal and bacterial taxa using Illumina MiSeq sequencing on branches of potted apple trees that had been inoculated with either water or a suspension of *N. ditissima* spores. We combined the knowledge on the metataxonomic profile with quantitative data on the *N. ditissima* branch colonisation (with relative abundances and absolute TaqMan qPCR concentrations) to identify taxa that show negative or positive correlations with *N. ditissima* DNA concentration. Secondly, we compared our fungal metataxonomic profile to the ITS amplicons of fungal isolates that had been tested for biocontrol potential in bioassays in a previous study. The aim was to possibly link fungal taxa with proven efficacy against the pathogen to the microbiome composition. The only ASVs showing a consistent negative correlation to relative and absolute *N. ditissima* abundance belonged to the bacterial genera *Kineococcus* and *Hymenobacter*. For fungal taxa only *N. ditissima* itself positively correlated to its increasing abundance, albeit only by rank and neither linearly nor beta binomially. Sequences belonging to the most promising antagonists from the study by Elena et al. (2022) could not be detected in the fungal microbiome profile at all. In addition, the combination of short reads length and high conservation within the chosen amplicon resulted in insufficient resolution to differentiate between a range of different efficacies of isolates belonging to the same genus (i.e. *Aureobasidium*).

## Introduction

European canker, caused by *Neonectria ditissima*, (syn. *Nectria galligena*; anamorph *Cylindrocarpon heteronema*) can lead to severe losses in apple production in the North-western regions of Europe as well as in Chile and New Zealand, where temperature is moderate, and rainfall is high throughout the year [1–3]. Infection occurs through different type of wounds such as pruning cuts [4], leaf scars [5] and fruit picking wounds [6]. The most important symptom is the development of cankers that can be formed in apple twigs, branches or/and the main stem, where it may lead the death of the whole tree [7, 8]

European canker management measures are not sufficiently effective. Soil conditions during the wet season grant only limited access to the orchards for the heavy machinery needed for fungicide sprays at leaf fall and the time window of application of fungicides is unfavourable due to their phytotoxic damage to the fruit surface (summarized in [9]). Nurseries can be an important source of infection since infected plant material can remain asymptomatic for months or even years and can arrive at the field apparently free of infection although it carries the pathogen [6, 8, 10]. Apple cultivars also differ in their levels of susceptibility to *N. ditissima*. Several studies have been performed to establish rankings of cultivar susceptibility [11, 12]. However, the understanding of the genetic basis of such resistance is still fragmented, hampering breeding for durable resistance [13]. Only a few studies have addressed the biological control of *N. ditissima* [14–17], none resulting in biocontrol products commercially available on the market to manage this disease. Recently advances have been booked on the biological control of *N. ditissima* by the work of Elena et al (2022) [18]. They isolated fungi at the boundary of canker and healthy tissue[19] and tested the resulting 520 fungal candidates for their efficacy of controlling of *N. ditissima* following a stepwise screening approach developed by Köhl et al. [19]*. Clonostachys rosea* isolates were the most promising antagonists against *N. ditissima*, by significantly reducing canker symptom severity and *N. ditissima* DNA concentration in the bioassays performed using detached ‘Elstar’ apple branches.

In addition to biological control as an environmentally friendly alternative, or besides to conventional disease management strategies, also the potential of the plant as a holobiont (the plant and its phytomicrobiome) is getting more in the focus of researchers. Members of the microbiome can provide wide array of direct or indirect biocontrol mechanisms protecting the plant from pathogens, facilitate uptake of essential nutrients and/or stimulate growth [20]. As the main routes of entry by *N. ditissima* are located on the above-ground parts of the apple tree (i.e. leaf scars, bark cracks etc) the microbial communities of the wood surface and internal tissues could play an important role in protecting the plant against infection. Metabarcoding of fungal and bacterial marker genes have been used previously to define the structure of apple tree endophyte communities associated with resistant/susceptible cultivars against *N. ditissima*, location and tissue type [21, 22]. It was suggested that bacterial and fungal species were enriched in resistant cultivars that are known to have antagonistic properties against other plant diseases. However, there was no direct link between the microbiome and the pathogen in these studies. To this end we combined a community analysis of bacterial 16S and fungal ITS barcodes with a study lead by Harteveld et al (2023) on colonization of apple branches by *N. ditissima* and the results on antagonistic fungi against *N. ditissima* isolated by Elena et al (2022) from infected apple branches [18, 23].

The aim of this current study is to combine the knowledge on the microbiome of the host plant in selected samples taken during the study of Harteveld et al. [23], using 16S and ITS metabarcoding with quantitative data on the *N. ditissima* branch colonisation to see if microbial components that show negative or positive correlations with *N. ditissima* concentrations can be identified. In a next step, ITS amplicons from the metabarcoding and those of the collection of potential antagonists against *N. ditissima* acquired in the study of Elena et al. (2022) were compared to evaluate if and how metabarcoding analysis can support decisions on selection of biological control agents [18].

## Material and Methods

### Experimental setup

The experiment using potted tress and the sample collection were described in detail in Harteveld et al. [23]. In short: 40 2-year-old potted trees of cultivars ‘Elstar’ (moderately resistant) and ‘Gala’ (susceptible) were used to study progression of *N. ditissima*. Half of the trees were inoculated with 100 macroconidia of *N. ditissima* per wound and half with sterile water as a control onto 5 fresh wounds per tree inflicted by cutting. Progression of *N. ditissima* was followed by applying a *N. ditissima*-specific TaqMan qPCR assay over a time of 8 weeks [18]. For the metabarcoding study 2 mm wood discs were cut from the site of inoculation at T0 (before inoculation), T1 (3 hours after inoculation) and T3 (4 weeks after inoculation).

### DNA isolation and PCR

A total of 184 wood discs from the wound surface were selected for DNA isolation, consisting of 124 samples treated with *N. ditissima* and 60 control samples from the two cultivars and three timepoints. Samples were lyophilized and homogenized by bead beating (6.35 mm RVS bead) with the Precellys (Bertin, Montigny-le-Bretonneux, France) for 2 x 15 sec at 6000 rpm with a 5 sec break. DNA was extracted with the MagAttract PowerSoil DNA extraction kit (Qiagen) according to the manufacturer’s protocol, DNA solution was diluted 1:1 and the fungal ITS2 region was amplified using primers ITS2KYO2F (5’-GATGAAGAACGYAGYRAA-3’) and ITS4R (5’-TCCTCCGCTTATTGATATGC-3’) (White et al. 1990) tagged with Illumina adapters in PCR reactions using a Q5® Hot Start High-Fidelity DNA Polymerase (New England Biolabs). The 25 amplification cycles were as follows: A denaturation step of 30 sec at 98°C followed by 25 amplification cycles of 30 sec 98°C, 30 sec 50°C, 30 sec 72°C and a final elongation step of 2 min at 72°C. The V4 region of the 16S SSU rRNA gene was amplified from undiluted samples in 35 cycles using primers 515F (5’-GTGCCAGCMGCCGCGGTAA-3’) and 806R (5’-GGACTACHVGGGTWTCTAAT-3’) also including Illumina adapters (Caporaso et al. 2011, PNAS). 5µM of mPNAs and pPNAs (PNAbio, USA) each were added to PCR mix. The PNA clamps mPNA and pPNA, block co-amplification of respectively plant mitochondrial and plastid targets in the samples [24] and depending on amount of bacterial target in the sample, will subsequently reduce presence of plant reads in sequencing data. The amplification protocol for 16S consisted of an initial denaturation at 98°C for 3 min, followed by amplification for 35 cycles at 95°C for 30 sec, 75°C for 10 sec for PNA annealing, 50°C for 30s for primer annealing and extension at 72°C for 30 sec, and a final extension of 1 min at 72 °C. For the ITS and 16S PCRs a dilution series (1□10^−1^-1□10^−3^) of ZymoBIOMICS Microbial Community DNA Standard (Zymo Research, Germany) was included in duplicate, as well as a negative extraction control. These samples were also included in the sequencing runs.

### Sequencing

Amplicons were purified and barcoded libraries were prepared according to the Illumina guidelines (Illumina, San Diego, CA, USA) and sequenced paired-end with 2×250 cycles for 16S V3-V4 libraries and 2127 cl:204300 cycles for ITS2 libraries on an Illumina MiSeq platform (Next Generation Sequencing Facilities, Wageningen University & Research, Wageningen, The Netherlands). A size selection step was included for the 16S amplicons before barcoding PCR in sequencing library preparation to remove amplicons smaller than 200 bp (primer dimers) with BluePippin (Sage Science, Beverly, MA, USA) pulsed-field electrophoresis on a 2% agarose gel. After sequencing, all reads were demultiplexed and quality checked before further bioinformatic processing. Raw data is available at NCBI under Bioproject ID PRJNA1036400.

### Sequencing data analysis

The sequences were processed using Qiime2 (version 2020-11, [25]) starting with the removal of primers and possible read-through using *cutadapt* ([26]). Reads smaller than 100bp were discarded. The *dada2* plugin [27] was used to trim the reads to a quality score of 10 and an expected error rate of 5%. The two 16S sets were then merged into a single abundance table. Taxonomy was assigned at 99% similarity with the UNITE database (version 8; https://unite.ut.ee) for the ITS Amplicon Sequence Variants (ASVs) and with the SILVA database (version 138; https://www.arb-silva.de) for the 16S ASVs. All ASVs were then filtered based on taxonomy: ASVs that were not classified as belonging to the kingdom ‘Fungi’ were removed from the ITS set. ASVs that could not be identified at the phylum level or were mitochondrial or chloroplast-derived were removed from the datasets, as well as ASVs with an abundance lower than 10 reads per sample and samples with less than 1000 reads in total.

Data were further analysed and visualized using RStudio (version 4.0.2). To investigate the original microbial community, the fungal and bacterial datasets were sub-set by “T0”. Cultivar-derived differences were investigated using the R package *MicrobiotaProcess* [28]. In a first step alpha indexes were calculated (*get_alphaindex*, Supplementary Figures 1A&B), followed by calculating the beta diversity to determine differences between microbiomes of the two cultivars at T0 (*get_pcoa*, distmethod=“bray”, method=“hellinger”).

The mean abundance of taxonomic classes across samples was calculated by first aggregating samples to a certain taxonomic rank (*tax_glom*, phyloseq), then converting counts to relative abundances (*transform_sample_counts*, phyloseq) and calculating rowMeans per taxon rank over all samples.

Correlation of the taxonomic composition between *N*. *ditissima* inoculation versus treatment with water was determined at T3 (no significant differences were found between categories at T0 and T1) for the two cultivars separately by sub-setting the both datasets by “T3”, converting the centered log ratio transformed data (microbiome::transform, *clr*) to a distance matrix (phyloseq::distance, “euclidean”) and doing a permutational multivariate analysis running the *adonis* function (vegan package [29]) on the ‘Treatment’ variable with 999 permutations to test for statistically significant differences. Of those results that differed significantly, similarity percentages were calculated using the *simper* function (vegan, permutations=100) showing to which extent ASVs contributed to the difference between the two groups (*N. ditissima*-treated vs H_2_O). To check for a correlation between components of the microbiome and *N. ditissima,* the *N. ditissima* DNA concentrations as obtained from the TaqMan qPCR assay and the relative abundances of the *N. ditissima* ASV were used separately. Samples that scored positive on *N. ditissima* presence were extracted and significant differential abundance of taxa along the gradient of *N. ditissima* concentration was determined by β-binomial regression (*corncob* [30]) at a false discovery rate of 10%. Sample T1_107_E_Nd_14_A was removed before the analysis as it contained 100% *N. ditissima*. For all analysis steps a p-value of 0.1 was accepted as significant. To analyze the relationship between the microbiome component *N. ditissima* and the *N. ditissima* concentrations as obtained from the TaqMan qPCR assay a separate linear regression and Spearman rank correlation were used.

### Identification of the isolates selected for the bioassays in planta

A total of 158 isolates were selected in the stepwise screening approach carried out by Elena et al. (2022) based on their spore production, their ecological characteristics for field applications, and the lack of environmental risks or pathogenicity known from literature for the selected species. Mycelia and spores or yeast cells of these isolates were used for isolation of genomic DNA. Total DNA was extracted and the primer set ITS1/ITS4 [31] was used for identification (DNA extraction, amplification and sequencing protocol are described in Elena *et al*. [18].

### [32]Antagonist database

To compare the 158 isolates from Elena et al. to the metataxonomic profiles obtained in this study we converted the ITS1/ITS4 sequences from these isolates into a blast database and used the sequences as queries for blastn alignment against the fungal metataxonomic reads (blast version 2.11.0+; [32]). Alignments with less than 100% sequence identity were checked for their taxonomic origin against the RefSeq database and only kept if they could be assigned to the same species.

### Efficacy test in bioassays in planta

The detailed description of the setup of the bioassays is described in Elena et al [18]. As a summary: the selected 158 isolates were tested in bioassays *in planta* using detached ‘Elstar’ apple branches. Artificial wounds on these branches were inoculated first with a spore suspension of *N. ditissima* and 24 hours later with a spore suspension of the corresponding isolate. Positive controls were wounds on branches only inoculated with *N. ditissima*. Inoculated branches were kept at 17 °C, 16 h light per day and RH>90% for four weeks. Thereafter, the ability of the isolates to prevent the growth and progress of *N. ditissima* along the branches was assessed by determining and comparing with the positive controls the symptom severity of the branches (ranging from 0: no symptoms to 4: severe canker) and the DNA concentration of *N. ditissima* determined by TaqMan qPCR assay [18] in discs taken from the inoculated branches, 0.5 cm distance from the inoculation site. *N. ditissima* DNA concentration quantified on the branches as pg/disk was transformed using log_10_ (x+1). Estimated marginal means (of 5 replicates) of log_10_ (x+1) obtained in the statistical analysis were back transformed. Percentage of reduction of *N. ditissima* DNA concentration on the branches was calculated from the back-transformed data, taking as reference the back-transformed data from the *N. ditissima* DNA concentration quantified in the positive controls per each bioassay. Canker symptom severity data was analysed with an ordinal mixed model to determine significant differences between the canker symptom severity on the branches inoculated with pathogen and candidate and the canker symptom severity of positive controls. A fitted linear mixed model was used to analyse significant differences between the DNA concentration of *N. ditissima* quantified on the branches inoculated with pathogen and candidate and the DNA concentration quantified on the positive controls per each bioassay. Additionally, the analysis of variance (ANOVA) of the linear mixed model was performed to identify the significant treatment effects.

## Results

### Sequencing stats

A total of 14,617,041 fungal ITS reads and 21,108,667 bacterial 16S reads were obtained from 191 samples. Due to technical problems one sample did not generate any reads. After filtering 12,274,171 ITS reads and 13,939,045 16S reads remained. Merging the paired end reads resulted in 7,140,970 ITS and 11,811,107 16S reads, representing 631 fungal ASVs and 2,115 bacterial ASVs. After filtering for low abundance (<10 reads), discarding chloroplast and mitochondrial sequences and ASVs that could not be assigned to a phylum, 349 fungal ASVs and 1154 bacterial ASVs remained. From these we excluded all samples that contained <1000 reads and removed all ASVs that were only found in these excluded samples resulting in 179 samples and a total of 323 AVS for ITS and 1079 ASVs in 184 samples for the 16S dataset.

### Independent molecular detection of N. ditissima using a TaqMan qPCR assay

The *N. ditissima* DNA concentration measured by the TaqMan qPCR assay increased at the wound surface of both cultivars from three hours (T1) after inoculation to 4 weeks (T3) after inoculation. The water-treated control samples did not show a significant quantitative increase over that timeframe (results relevant for this study are summarized in Fig. 1).

**Figure 1:**
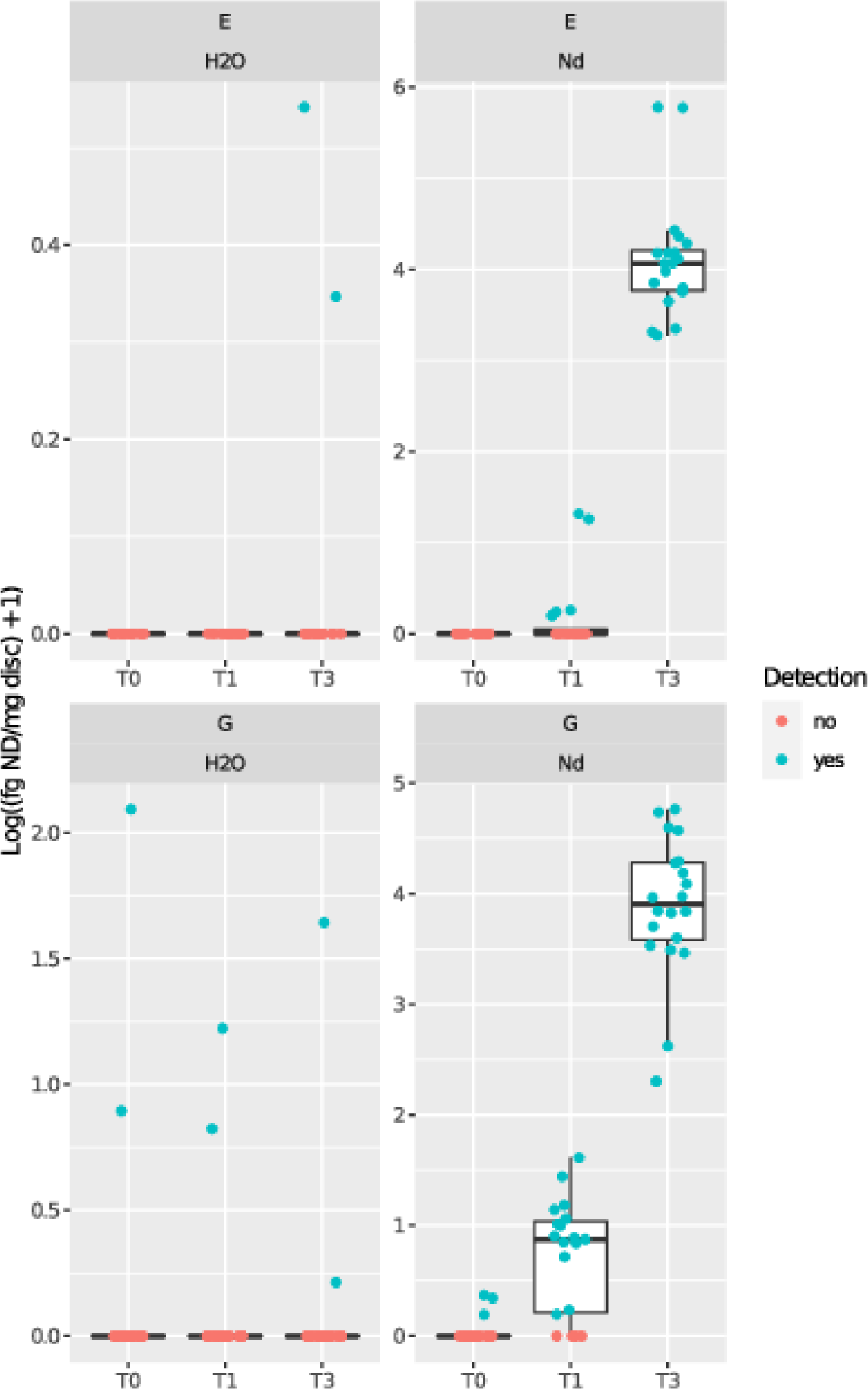
Concentration of *N. ditissima* DNA in control (H2O) and inoculated (Nd) wood discs over time T0 (after inoculation), T1 (3h post inoculation), T3 (4 weeks post inoculation) in apple cultivars ‘Gala’ (G) and ‘Elstar’ (E).

### Microbiome shifts over time

#### 16S

The bacterial twig microbiome composition at the starting situation (T0) showed that although the relative abundance of taxa varied in different trees of the same cultivar, the core microbiome they shared was very similar (Fig. 1 A&B). This was confirmed by calculating the beta-diversity showing a high degree of overlap between samples of the two cultivars (Supplementary Figure 2A). In ‘Elstar’ the most abundant bacterial orders (expressed as mean relative abundance over all samples) were Burkholderiales (22.0%), Sphingomonadales (19.6%), Rhizobiales (9.7%), Propionibacteriales (9.1%), Micrococcales (7.0%) and Pseudomonadales (6.8%). In the susceptible cultivar ‘Gala’ the same orders were found, but the Sphingomonadales represented the most abundant group with 29.0%, followed by Burkholderiales (15.2%), Rhizobiales (11.3%), Micrococcales (9.0%), Propionibacteriales (8.0%) and Pseudomonadales (6.9%) (Supplementary Table 1).

Three hours after inoculation (T1) with either inoculation with a *N. ditissima* spore suspension or water the bacterial microbiome changed significantly (Fig. 1 A&B). The relative abundance of the order Burkholderiales increased to 79.0% in ‘Elstar’ and 58.0% in ‘Gala’. Whereas at T0 the Burkholderiales consisted mostly of members of the families *Comamonadaceae* and *Oxalobacteraceae* (and *Neisseriaceae* in ‘Gala’) (Supplementary table 2A&B). The steep increase in the Burkholderiales at T1 was due to a single ASV belonging to the genus *Ralstonia.* Its relative abundance ranged from 42.9-92.0% in ‘Elstar’ and 0.4-86.2% in ‘Gala’ disregarding the treatment. This ASV was present at T0 in ‘Elstar’ in a single sample (5.3%) only and could not be detected in ‘Gala’ samples at T0. Four weeks after inoculation (T3) this specific *Ralstonia* ASV was detected in neither ‘Gala’ nor ‘Elstar’ anymore. The bacterial population resembled the situation at T0. At T3 ‘Gala’ and ‘Elstar’ microbiomes were comparable, with Sphingomonadales (40.1% and 37.6%) being the most dominant order in both cultivars (Supplementary Table 1). The relative abundance of the order Micrococcales had increased by threefold compared to T0 with *Curtobacterium* and *Frondihabitans* being the most abundant genera.

#### ITS

The original fungal microbiome sampled before inoculation (T0) did show differences between the two cultivars (Supplementary Figure 2B) and was also more stable over time (Figure 2 A&B). The most abundant genera detected in ‘Elstar’ before inoculation were *Vishniacozyma* (30.6%), *Alternaria* (16.3%), *Cladosporium* (9.7%), *Filobasidium* (9.3%), *Aureobasidium* (8.9%) and *Erythrobasidium* (6.4%). In ‘Gala’ also *Vishniacozyma* (29.2%), *Cladosporium* (15.2%) and *Alternaria* (14.3%) were the three most abundant genera, followed by *Didymella* (9.8%), *Aureobasidium* (6.6%) and *Cadophora* (5.2%) (Supplementary Table 1).

**Figure 2:**
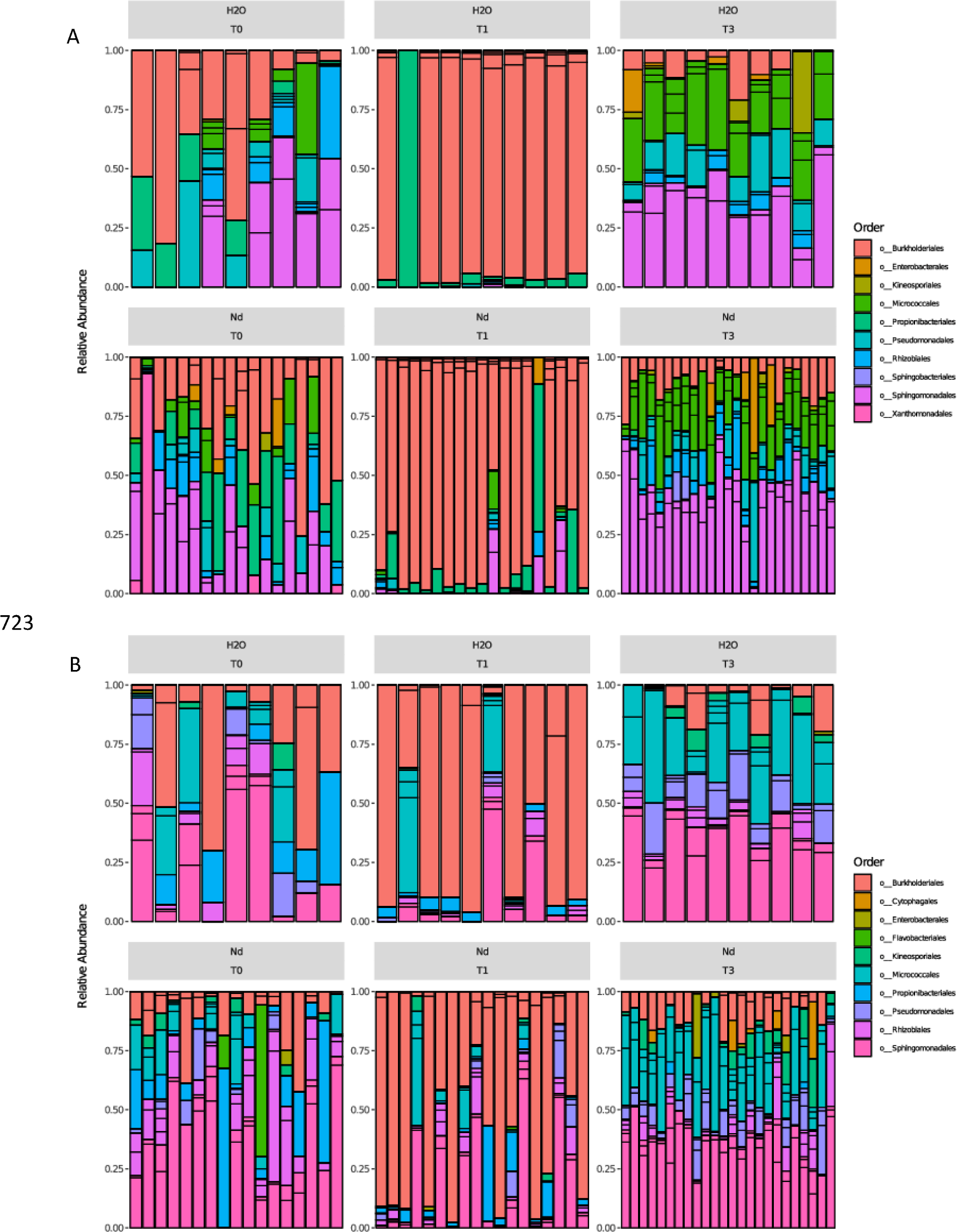
Relative abundances of the 25 most abundant bacterial 16S ASVs in wood discs of apple cultivars ‘Elstar’ (A) and ‘Gala’ (B) in treated (Nd) and control samples (H2O) over the time of the experiment. T0 (after inoculation), T1 (3h post inoculation), T3 (4 weeks post inoculation). ASVs were grouped at the level of order. Bars represent different ASVs within a taxonomic group.

**Figure 3:**
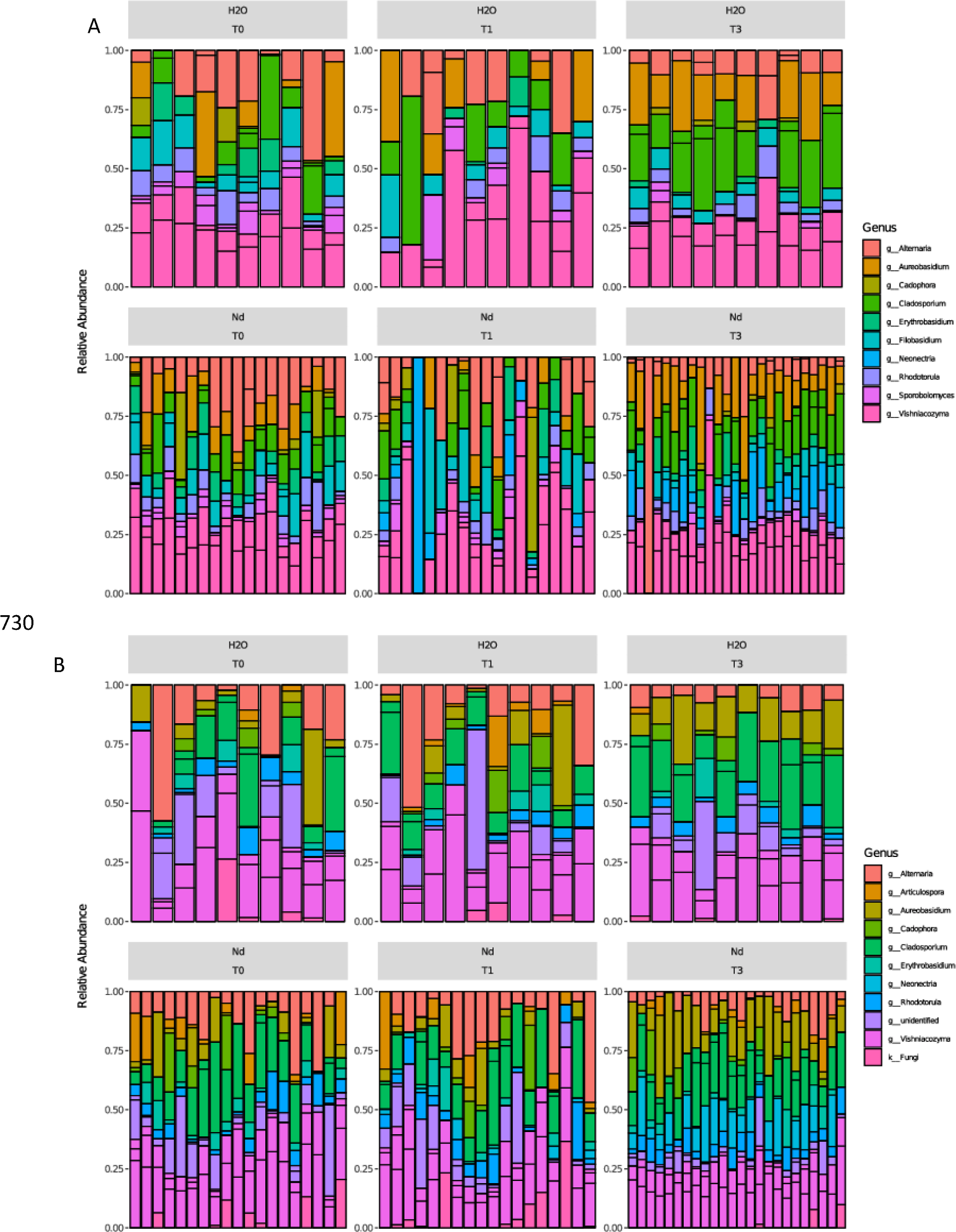
Relative abundances of the 15 most abundant fungal ITS ASVs in wood discs of apple cultivars ‘Elstar’ (A) and ‘Gala’ (B) in treated (Nd) and control samples (H2O) over the time of the experiment. T0 (after inoculation), T1 (3h post inoculation), T3 (4 weeks post inoculation). ASVs were grouped at the level of genus. Bars represent different ASVs within a taxonomic group.

The most dominant taxa stayed the same three hours (T1) and four weeks (T3) after inoculation, with ‘Elstar’ differing from ‘Gala’ in containing relatively higher numbers of *Filobasidium* and ‘Gala’ having a higher relative abundance of *Didymella* (Supplementary Table 1). In samples treated with the *N. ditissima* suspension, the average relative abundance of ASVs that could be assigned to the genus *Neonectria* increased over time from absent in ‘Elstar’ and ‘Gala’ at T=0 to 6.9% in ‘Elstar’ and 7.33% in ‘Gala’ until 4 weeks after inoculation (T3).

### Correlation of microbiome data with relative and absolute Neonectria concentrations

Due to this difference in the pathogens’ DNA concentrations between the *N. ditissima*-treated and the control samples at T3, it was tested whether the treatment itself did have a significant effect on the microbial composition at T3 in the two cultivars. For the bacterial dataset this was not the case in ‘Gala’ (R^2^=0.03573, p=0.187), but in ‘Elstar’ significant differences in the microbiome compositions between the *N. ditissima*-treated samples and the control samples were found, although the effect sizes were small (R^2^=0.03986, p=0.085). Also, the fungal dataset did show significant differences between the treatments at T3 in both cultivars (R^2^=0.0661 (‘Elstar’), 0.05942 (‘Gala’), p<0.001). In the datasets showing significant differences the contribution of each ASV to the dissimilarity between the two treatment groups was calculated (Supplementary Table 6). In the bacterial ‘Elstar’ dataset ASVs that had the highest contribution to the overall dissimilarity were two members of the family *Microbacteriaceae* (5.7%, p=0.099 and 2.74%, p=0.0198) and a member of the genus *Kineococcus* (2.4%, p=0.0198). In the fungal ‘Elstar’ dataset the differences were caused by members of the genus *Cladosporium* (4.9%, p=0.0198) and *N. ditissima* (4.8%, p=0.0396). In the ‘Gala’ ITS dataset the highest contribution to the overall dissimilarity was attributed to *N. ditissima* (3.7%, p=0.0297), *Cadophora* (2.2%, p=0.0396) and *Didymella* (1.5%, p= 0.0099). As could be expected, the relative abundance of *N. ditissima* was higher in the samples treated with the pathogen. The relative abundance of the of the other ASVs was higher in the water-treated control. The relative abundances of the differential abundant ASVs were plotted against the relative abundance of *N. ditissima* (Supplementary Figures 3A and 3B) as well as the absolute concentration of *N. ditissima* DNA (Supplementary Figures 4A and 4B). The only ASVs showing a consistent negative correlation to *N. ditissima* abundance with both methods belonged to *Kineococcus* and *Hymenobacter*. The latter was only present in four samples.

The treatment category is a non-numeric, qualitative parameter that does not take variability of *N. ditissima* concentrations between and within categories into account. To make more accurate correlations, microbial compositions had to be linked to absolute *N. ditissima* abundances. Due to the TaqMan results published in Harteveld et al. (2023) [23], absolute abundances of *N. ditissima* were available for each sample. In addition, relative abundance of the pathogen per sample could be derived from the microbiome analysis. In a first step we tested whether the two measures of pathogen presence were comparable. Associations between the DNA concentrations of *N. ditissima* measured by TaqMan qPCR and the relative abundance of *N. ditissima* ASVs in the samples did show similarity (Supplementary Figure 4B) but were not found to be significant using beta-binomial regression (p=0.9). A linear regression with permutation of this same data also found no correlation between the two (p=0.36). Using a rank correlation, a significant nonlinear correlation with each dataset was found (p=2.2□10^−16^). Correlations between the abundance of the pathogen and the residual bacterial and fungal microbiome were done using both methods separately.

To test whether certain components of the microbial communities were either positively or negatively correlated with relative abundances of *N. ditissima*, multiple beta-binomial regression was performed. In all samples containing ASVs that could be assigned to *N. ditissima*, the relative abundance of *N. ditissima* was taken as an anker to determine which ASVs increased or decreased along the gradient of *N. ditissima* concentrations. This resulted in mostly bacterial ASVs shown in Table 1. ASVs that decreased in both cultivars while *N. ditissima* increased belonged to *Hymenobacter* and to the *Methylobacterium-Methylorubrum* group as well as a member of the family *Sphingomonadaceae* (possibly *Novosphingobium*). In ‘Gala’ also ASVs of the genera *Cutibacterium*, *Kineococcus, Mucilaginibacter* and the fungal genus *Cadophora* decreased with increasing *N. ditissima* concentrations. Few ASVs also increased with a higher relative abundance of *N. ditissima*, including *Neonectria* itself, one ASV of the fungal genus *Vishniacozyma*, and unknown basidiomycete and an ASV belonging to the genus *Pseudomonas*.

**Table 1:** Fold change increases of the relative abundance of bacterial and fungal ASVs per 1% increase in the relative abundance of *N. ditissima* at T3 (4 weeks post inoculation). P-values are corrected for multiple testing.

| Time | Cultivar | Lowest tax level | ASV | Highest identified tax level | Slope<sup>a</sup> | P-value |
| --- | --- | --- | --- | --- | --- | --- |
| T3 | Gala | Bacteria | 5a7b179b1b45f0fe2282f260bf073f60 | *Cutibacteria* | -63 | 0.01 |
| T3 | Gala | Bacteria | 23620e81538019f85c727ec1791df244 | *Mucilaginibacter* | -38 | 0.03 |
| T3 | Gala | Bacteria | 4a6086b0d2aac9c15dcc4c55f3c38f79 | *Hymenobacter* | -30 | 0.01 |
| T3 | Elstar | Bacteria | 4a6086b0d2aac9c15dcc4c55f3c38f79 | *Hymenobacter* | -16 | 0.05 |
| T3 | Gala | Bacteria | 8b7ed699511c4ce2d3e390b6d3ffab03 | *Sphingomonadaceae* | -15 | 0.08 |
| T3 | Gala | Fungi | 385465619e3674e430081def76e0daae | *Cadophora* | -14 | 0.01 |
| T3 | Elstar | Bacteria | 8b7ed699511c4ce2d3e390b6d3ffab03 | *Sphingomonadaceae* | -12.5 | 0.06 |
| T3 | Gala | Bacteria | 374b724806294b861c4ed5cfb74e2f31 | *Methylobacterium-Methylorubrum* | -10 | 0.02 |
| T3 | Gala | Bacteria | b1dde0ed7ab1f036845b5643cfaa699e | *Kineococcus* | -9 | 0.08 |
| T3 | Elstar | Bacteria | 374b724806294b861c4ed5cfb74e2f31 | *Methylobacterium-Methylorubrum* | -7.5 | 0.05 |
| T3 | Gala | Fungi | 80903b10b9a6b7af1e25c8a08ee5d4ff | *Vishniacozyma* | 1 | 0.07 |
| T3 | Elstar | Fungi | 04be2f423b3a9a05b222355a42d59f39 | *Neonectria* | 6 | 0 |
| T3 | Gala | Bacteria | b4289ab0a9a00034c1fbce477b2c4ded | *Pseudomonas* | 6 | 0.03 |
| T3 | Gala | Fungi | 04be2f423b3a9a05b222355a42d59f39 | *Neonectria* | 16 | 0 |
| T3 | Elstar | Fungi | 2dae6151605d88893bfae16aefeb4ad1 | *Basidiomycota* | 54 | 0 |

<sup>a</sup> increase of ASV relative to 1% increase to *Neonectria ditissima*

Then all samples in which the concentration of *N. ditissima* determined by TaqMan qPCR was above the detection limit were subjected to the same analysis. Again, the bacterial genera’s relative abundance that decreased with higher *N. ditissima* concentrations were dominant (Table 2). For ‘Gala’ similar correlations were found for the ASVs of *Hymenobacter*, *Kineococcus* and *Methylobacterium-Methylorubrum* as were shown previously. In addition, *Sphingomonas* and a member of the *Microbacteriaceae* (*Frondihabitans*) also decreased. In ‘Elstar’ the only genus that decreased slightly when *N. ditissima* concentrations increased was *Massilia*. The only fungal genus increasing relative to increasing *N. ditissima* concentrations in ‘Gala’ was *Neonectria* itself.

**Table 2:** Fold change increases of the relative abundance of bacterial and fungal ASVs per fg/ul increase of *N. ditissima* at T3 (4 weeks post inoculation). P-values are corrected for multiple testing.

| Time | Cultivar | Lowest Taxum | ASV | Highest identified Taxum | Slope<sup>b</sup> | P-value |
| --- | --- | --- | --- | --- | --- | --- |
| T3 | Gala | Bacteria | 4a6086b0d2aac9c15dcc4c55f3c38f79 | *Hymenobacter* | -0.00125 | 0 |
| T3 | Gala | Bacteria | b1dde0ed7ab1f036845b5643cfaa699e | *Kineococcus* | -0.0006 | 0.01 |
| T3 | Gala | Bacteria | 374b724806294b861c4ed5cfb74e2f31 | *Methylobacterium-Methylorubrum* | -0.0006 | 0 |
| T3 | Gala | Bacteria | 9ac3bb1d7dedb08a012692a6f536b5af | *Sphingomonas* | -0.0002 | 0.03 |
| T3 | Gala | Bacteria | f32490b8845851fe9be12251653f85f3 | *Microbacteriaceae* | -0.00012 | 0.02 |
| T3 | Elstar | Bacteria | ca6c2b3b469c08212142e8821c65882a | *Massilia* | -1.10E-05 | 0.03 |
| T3 | Gala | Bacteria | 497a26126c9e726c768bdec79e8c4e9b | *Curtobacterium* | 0.00013 | 0.03 |
| T3 | Gala | Fungi | 04be2f423b3a9a05b222355a42d59f39 | *Neonectria* | 0.00023 | 0 |
| T3 | Gala | Bacteria | b4289ab0a9a00034c1fbce477b2c4ded | *Pseudomonas* | 0.00025 | 0.03 |

<sup>b</sup>relative change of ASV per fg/ul increase of *Neonectria ditissima*

The corresponding ASVs and their relative counts were extracted and plotted to show their overall distribution in the samples (Supplementary Figure 3A&B for the relative data and 4A&B for absolute data). Especially *Cutibacterium, Mucilaginibacter, Hymenobacter* and the unknown basidiomycete had low relative abundances and/or were only present in a few samples.

### Extraction of antagonist sequences from microbiome sequences

The ITS amplicons of the selected 158 isolates by Elena et al. [18] were used to extract the closest hits from the fungal microbiome dataset using *blastn* with the aim of ranking the detected isolate sequences in the microbiome according to their efficacy in the bioassays. This would have allowed us to do a network analysis to get more insight into which taxa co-occur with antagonists and how that relates to *N. ditissima* abundance. However, only 18 ASVs could be matched to the isolates’ ITS sequences, 17 of which with a similarity >98%. This resulted in three ASVs matching 50 *Cladosporium* isolates with 100% similarity, a single ASV matching all 54 *Aureobasidium* isolates, a single ASV matching 15 *Didymella* isolates, 5 ASVs matching 23 isolates of different *Vishniacozyma* species (all 100% similarity), two ASVs matching the two *Sporidiobolus* sp. isolates in the collection, one ASV matching *Papiliotrema* sp., one matching *Gabarnaudia betae* (99.55%) and three ASVs matching the isolate of *Cystobasidium pinicola* between 98.55% and 99.71% (Supplementary table 7). Sequences belonging to the most promising antagonists, could not be detected in the fungal microbiome profile, nor was the taxonomic resolution sufficient to differentiate between efficacies of isolates belonging to the same genus. Therefore, no in-depth analysis of networks was performed. Nevertheless, we proceeded with the overlapping ITS sequences and plotted the relative abundances of the ASVs at T3, contrasting them on the treatment category showed the high variability of their relative abundance in samples in general (Fig. 4). Most of them did not show a significant difference between the pathogen inoculated samples and the control. A genus that was found to be significantly different (p<0.05) between treatment and control was *Didymella*. This difference was however restricted to ‘Gala’, where this ASV was found in three control samples and not in the treated samples. Also, a ASVs belonging to *Cladosporium* was relatively more abundant in control samples (p<0.1). These were the same ASVs that were found to be the main contributors to dissimilarity between the treatment categories mentioned previously. Two ASVs also did show significantly higher numbers (p<0.1) in the inoculated samples these being one of the ASVs belonging to *Sporobolomyces* and one belonging to *Cystobasidium*. However, none of the extracted sequences did show a significant correlation with any of the two quantitative *N. ditissima* values (Table 1 and 2).

**Figure 4:**
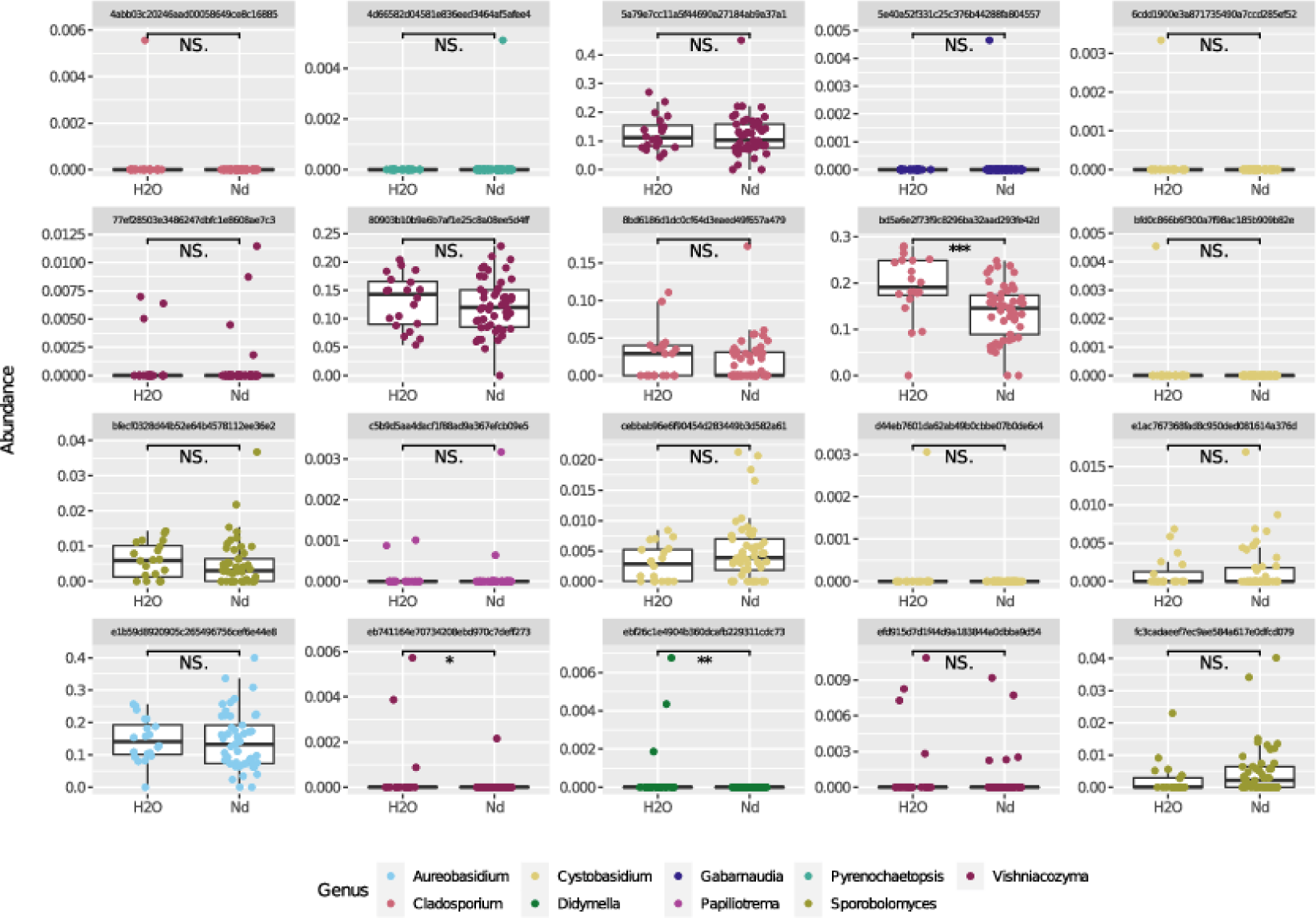
Relative abundance of ASVs matching the ITS sequences of isolates described in Elena et al. (2022) in water (H2O) or *N. ditissima* (Nd) inoculated wood discs.

## Discussion

The overall structure of the bacterial and fungal microbial community found in this study resembled that previously described in apple wood tissue as well as apple fruit in other metabarcoding studies [22, 33, 34]. In addition, taxa carrying these representative fungal sequences could also be isolated from symptomatic apple branches as has been described recently by Elena et al. (2022) showing that they are indeed active inhabitants of the above ground microbial apple community [18].

It has been hypothesized previously that certain components of the endogenous microbial community could contribute to susceptibility [35, 36] or resistance [37–39] of a plant to certain pathogens and/or severity of disease symptoms. Also, for European canker different levels of canker susceptibility have been linked to certain taxa when comparing metataxonomic profiles of apple cultivars with a varying degree of resistance to the disease [22]. This included the bacterial genera *Sphingomonas*, *Curtobacterium, Massilia, Hymenobacter, Frondihabitans, Pseudomonas* and *Methylobacter* as well as the fungal genera *Filobasidium*, *Vishniacozyma*, *Dioszegia* and *Gelidatrema* to be less abundant in resistant genotypes. The fungal genera *Aureobasidium, Rhodotorula, Stemphylium* and *Kalmanozyma* were relatively more represented in resistant cultivars [22]. Although contrasting resistant and susceptible cultivars is an interesting first step to describe differences between apple genotypes, extrapolation of these findings to identify microbes potentially involved in disease suppression is difficult. This is due to several reasons: the microbiome is dynamic depending on biotic and abiotic characteristics of the host plant and the environment, but also on the site of sampling (stem, leaf, root) [40]. Also, the identified microbiome may be genotype-specific, linking this to the resistance against a single pathogen is at least incomplete, because a gradient of resistances against multiple pathogens can exist in the same genotype and findings are therefore not exclusive to one pathogen. This problem can be minimized by choosing larger pools of resistant and susceptible cultivars against the pathogen in question, like done by Olivieri et al [22]. Still, it must be considered that there is not always a clear consensus about different degrees of resistance. Because the resistance criterion used in each study is different, correlations are less robust [4, 11, 12]. In addition, microbiomes are shaped by their exposure to the pathogen in question. It has been shown previously that the introduction of the pathogen triggers microbiome-mediated disease suppression [41]. This is an important factor to consider when screening microbiomes for components that help alleviate disease development.

The present study included the direct link to pathogen exposure by artificial inoculation of trees, the quantification of the pathogen and following the microbiome development along with the progression of the pathogen over time. This offered the possibility to (i) link microbial taxa to the treatment as well as (ii) the actual concentration or relative abundance of *N. ditissima* in each sample. Comparing the microbial profiles of pathogen-inoculated samples and the water control four weeks after the treatment, a time after which a clear difference in *N. ditissima* concentration could be measured by TaqMan qPCR between these two categories, resulted in very low effect sizes. Only 3.98% (16S, ‘Elstar’) to 6.61% (ITS, ‘Elstar’) of the variation in distances is explained by the grouping. This could be caused by the generally very high variation among the replicates. In addition, *N. ditissima* also occurred naturally in several control samples making the contrasting less reliable. The contribution of specific ASVs to the overall dissimilarity of *N. ditissima*-treated samples was also low. *N. ditissima* which was initially inoculated with approximately 100 spores per wound and could be detected by TaqMan qPCR in all treated samples at T3 accounted for only 4.82% or 3.71% of the differences in the ‘Elstar’ and ‘Gala’ microbiome respectively. The higher relative abundance of *N. ditissima* in pathogen-treated samples was to be expected. Other dissimilarities were explained by higher relative abundances of a few bacterial and fungal ASVs in the water-treated controls. This included two ASVs assigned to the *Microbacteriaceae*, *Kineococcus* and the fungal genera *Cladosporium, Cadophora and Didymella*. Most of these taxa have been described as common inhabitants of the phyllosphere in many plants before and have representatives with an endophytic lifestyle [42–45]. *Cadophora* is known as an inhabitant of the plants’ rhizosphere [46] and well equipped to thrive as endophyte in wood tissue where it can cause soft rot [47, 48]. Non-pathogenic isolates have however also been attributed with antagonistic characteristics and shown to suppress several pathogens in tomato *in vitro*. However, these observations could not be translated to *in planta* assays showing the complexity of interactions when adding the plant as a variable [49].

Due to the high variation in the concentration of *N. ditissima* at T3 and the fact that the pathogen was naturally present in a few control samples, limiting the analysis to comparing the treatment categories would have been insufficient. The exposure to the pathogen was not equally distributed across the samples and categories. Therefore, it was also investigated how the microbiome composition was build up across a gradient of *N. ditissima* concentrations. Here we could not identify any fungal taxon that correlated with the combined absolute and relative pathogen concentrations either positively or negatively, except for *N. ditissima* itself. Also, among the bacterial taxa only two ASVs shared a negative correlation with absolute and relative *N. ditissima* concentrations. These ASVs belonged to the genera *Kineococcus* and *Hymenobacter*. The *Hymenobacter* ASV was only present in 4 samples and therefore not considered further. *Kineococcus* is a genus in the phylum of Actinobacteria and has been isolated in very diverse environments including desert soils [50], marine sediments [51], xylem sap and wood chips [52]. Several isolates are known for their tolerance against γ-radiation, strong oxidants and metals such as copper and zinc as well as their production of indole acetic acid, a plant hormone of the auxin class, inducing plant growth [53, 54].

Although no fungal ASVs could be identified in significantly higher abundances when *N. ditissima* numbers were low, we still compared these metataxonomic profiles with the database of isolated potential antagonists selected in the stepwise screening approach performed by Elena *et al.* (2022) [18]. The reasoning behind this was that if promising antagonists already tested *in planta* could be detected in our metataxonomic profiles, we could possibly identify factors, such as networks of microorganisms they cooccur with, that shape the antagonist selection. Unfortunately, the metataxonomic profiles did not show overlap with the ones that came out best in efficacy trials of Elena et al. (2022) [18]. On the contrary *Clonostachys rosea,* the species with the most promising antagonistic effect, was not found at all. The fact that we could not retrieve any *C. rosea* can be explained by the different nature of the samples in combination with its potential mode of action. Whereas the samples for metataxonomic profiling were taken from apple trees four weeks after being artificially inoculated with *N. ditissima*, Elena et al. (2022) retrieved the isolates from trees showing well-established cankers, at the boundary between healthy and cankered tissue [18]. The composition of the microbiome is probably not the same, although overlap can be assumed, as the most abundant fungal ASVs in the metataxonomic profile resembled, the most abundant taxonomic groups gained by the cultivation-dependent approach by Elena et al. [18]. The biological control mechanism of *C. rosea* against pathogens is primarily attributed to the activation of multiple mechanisms such as secretion of cell□wall□degrading enzymes, production of antifungal secondary metabolites, such as antibiotics and toxins, and induction of plant defence systems [55–58]. Modes of action that require a direct contact between pathogen and antagonist and can be considered curative, show different dynamics and spatial-temporal development than preventive strategies. This shows that different modes of action of a beneficial community component can result in different correlations. [59, 60]. In some cases, it is necessary that the necrotrophic plant pathogens invade the tissue of host plants first and use the available nutrients as primary colonizers. This would result in positive correlations with the pathogens concentration. Once necrosis has been induced by the pathogen, non-pathogenic microorganisms with saprophytic lifestyle can also colonise necrotic tissues and play an important role in competitive substrate colonisation [59]. Accordingly, in tissues with higher concentrations of *N. ditissima,* potential antagonists might be present. In addition, a major drawback of short-read amplicon sequencing was confronted: insufficient resolution due to the short amplicon. For example, the *Aureobasidium* isolates tested by Elena et al. (2022) [18] had different efficacies against *N. ditissima* in bioassays but could not be distinguished on amplicon level, as *Aureobasidium* was condensed in a single ASV due its identical sequence in all species along the length of the amplicon. This made a differentiation between promising and unsuitable antagonists impossible.

This shows that correlations of the metataxonomic profile with either the treatment or with the concentration of *N. ditissima* in our samples are a first step to understand the population dynamics in the initial stages of disease development, their value in the identification of key microbial players in disease suppression and antagonist selection seems to be overrated. It does not consider the different ecology of antagonists (different modes of action) resulting in contrasting correlations with the pathogen, the variation in efficacies within the same genus or even species that cannot be distinguished and the changes in the microbiome composition, especially upon disturbances (i.e. cutting and inoculation in this case) in less diverse samples were short-term temporal shifts could easily lead to false positive or false negative correlations.

This study shows that a cautious interpretation of metataxonomic profiles and critical evaluation of its additional value to the selection process of biocontrol agents is recommended. To limit the mentioned drawbacks we advise to include a reasonable number of replicates to minimize the effect of a high variation between samples having received the same treatment, considering spatial-temporal variation (by including a time series), sufficient taxonomic resolution by switching to (synthetic) long-read sequencing [61] and possibly adding a quantitative dimension to bypass the compositional nature of the dataset and increasing the statistical power of biologically relevant interactions [62].

## Acknowledgements

This research was partly funded by the Dutch Ministry of Agriculture, Nature and Food Quality and a consortium consisting of 10 companies lead by the Dutch Fruit Growers Organisation (NFO) and was part of the project KV 1605-033 ‘PPS Integrale ketenaanpak vruchtboomkanker’. The authors would like to thank Dalphy Harteveld, Engelien Kerkhof, Ilse Houwers, Jan Cordewener and Elio Schijlen for their technical assistance (field trials, DNA isolation and sequencing) and Dennis te Beest for advice on statistical methods.

## Notes

https://www.ncbi.nlm.nih.gov/bioproject/?term=PRJNA1036400

### Competing Interest Statement

The authors have declared no competing interest.

