## Supplementary figures and images for "From amplicons to strains: The limitations of metabarcoding as criterium in the selection process of biocontrol strains against the pome fruit pathogen *Neonectria ditissima*"

### Supplementary_Fig1A_alpha_index_T0_16S.pdf

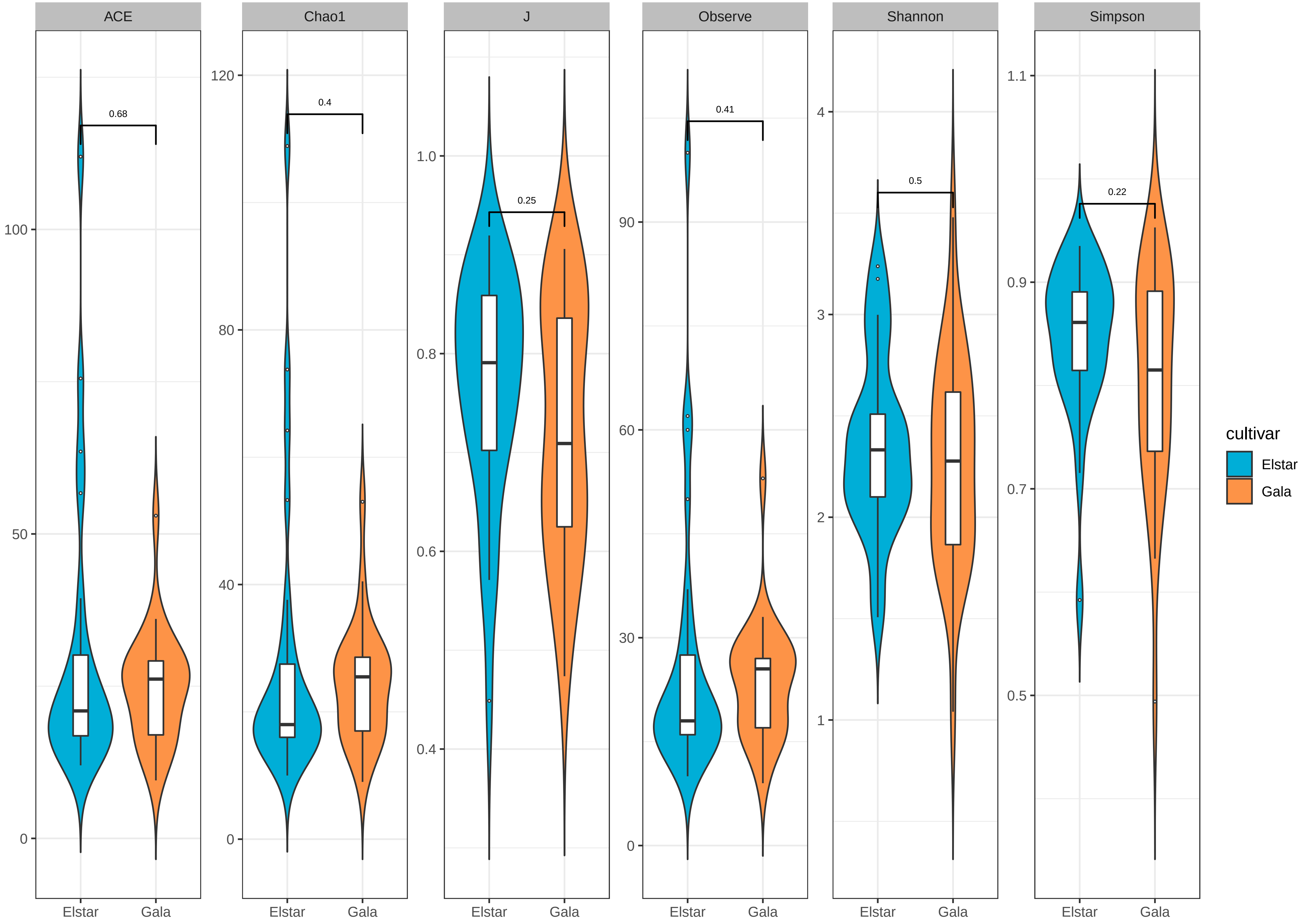

### Supplementary_Fig1B_alpha_index_T0_ITS.pdf

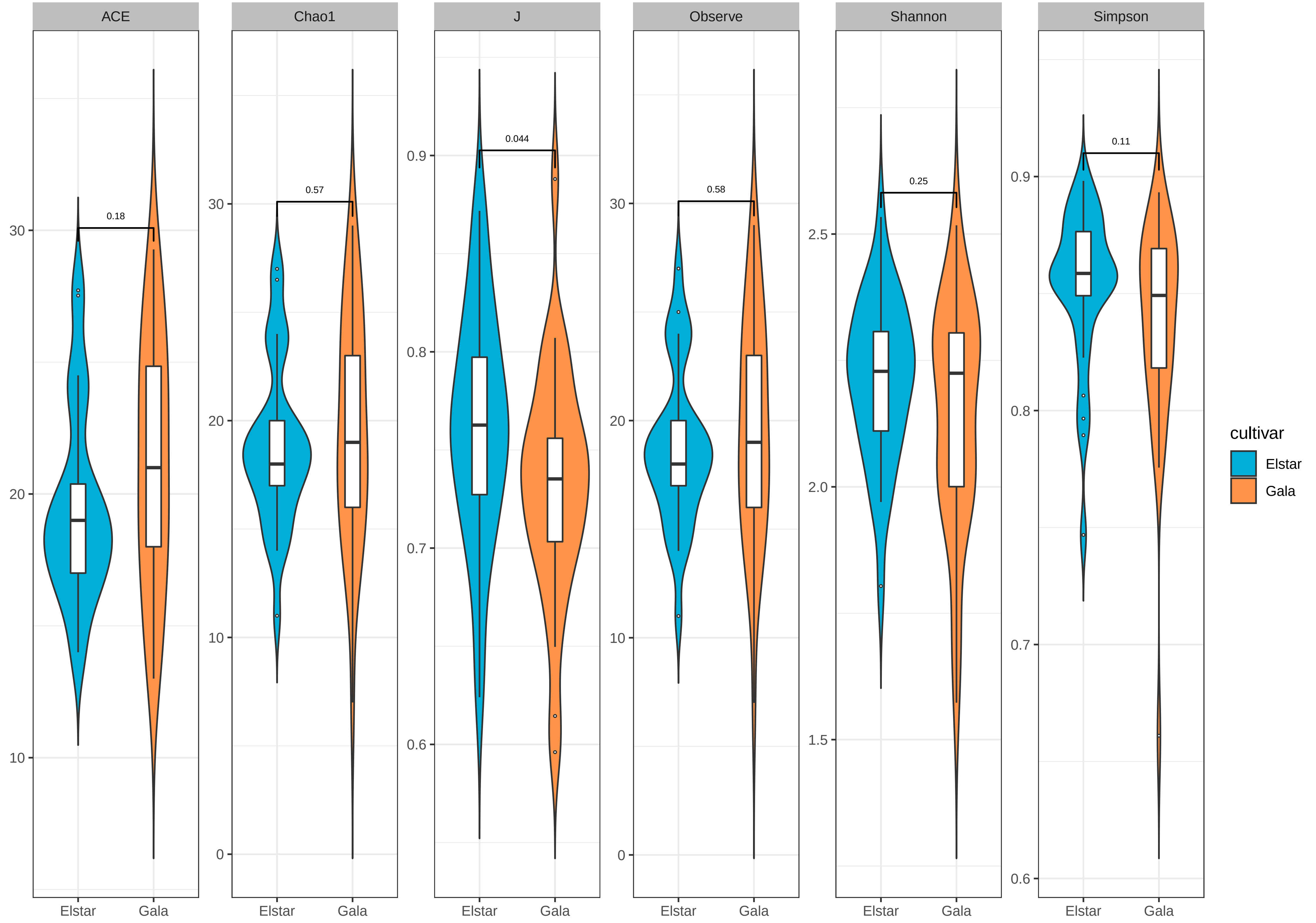

### Supplementary_Fig2A_pcoa_16S_T0.pdf

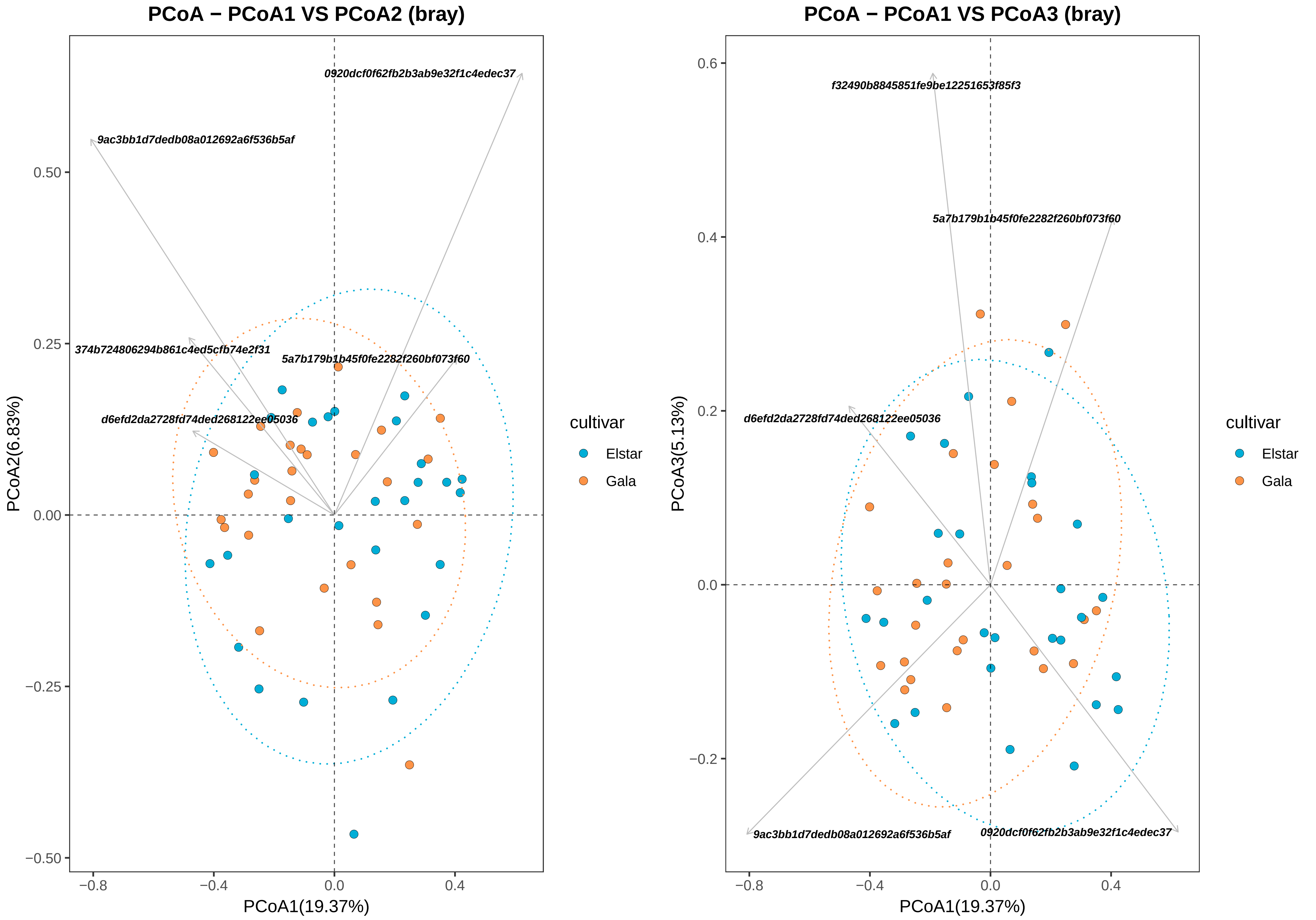

### Supplementary_Fig2B_pcoa_ITS_T0.pdf

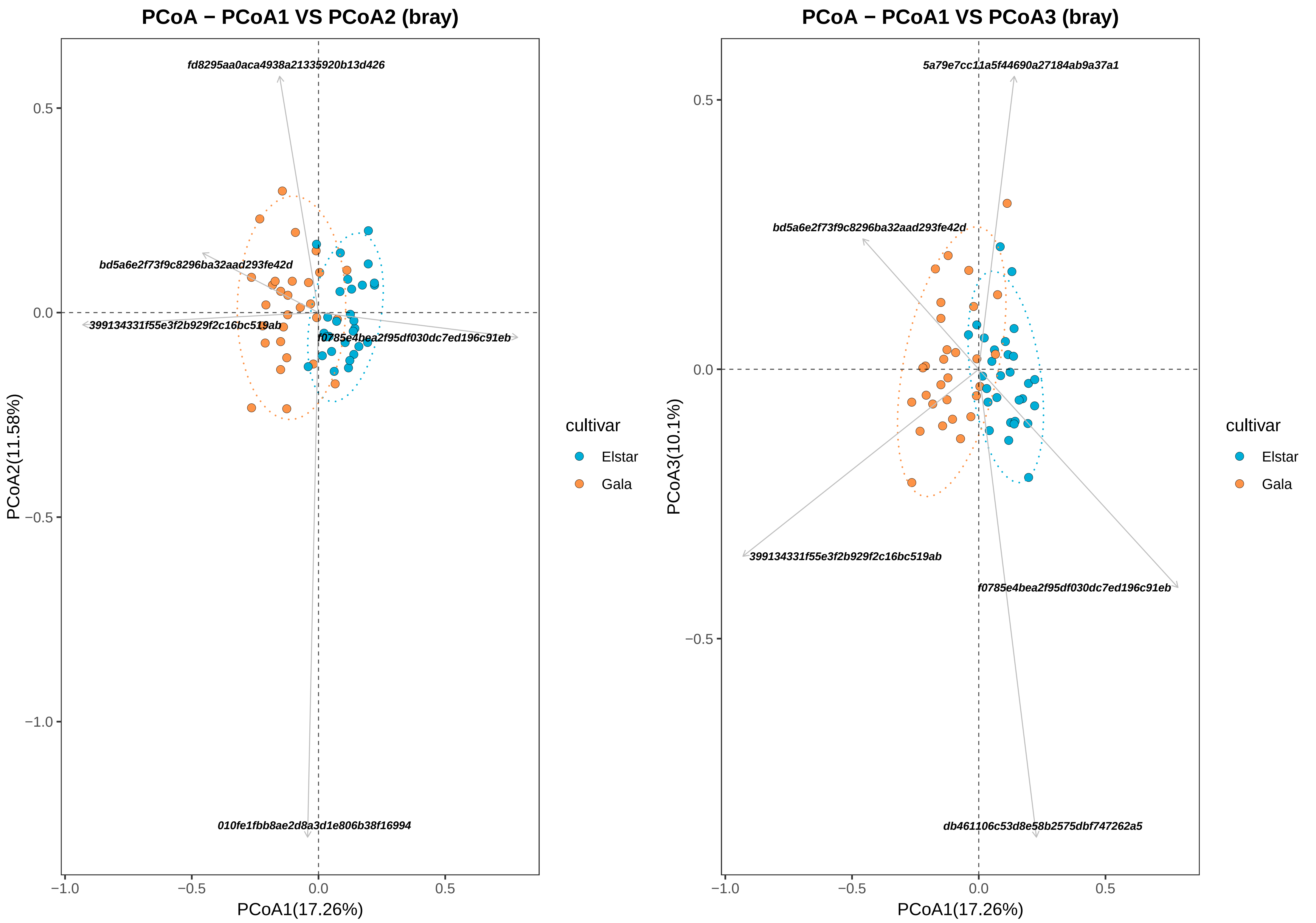

### Supplementary_Fig3A_rel_16S.pdf

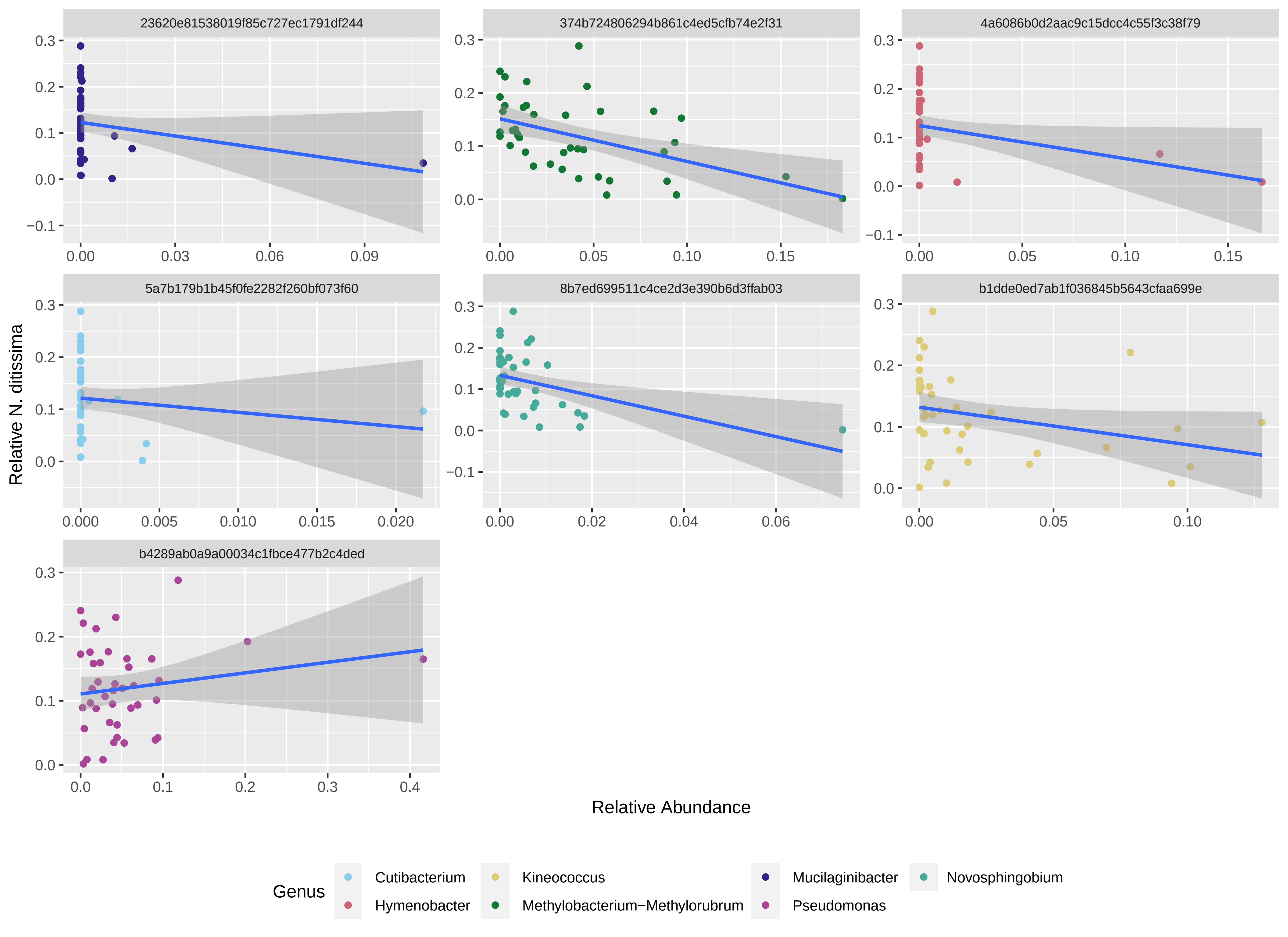

### Supplementary_Fig3B_rel_ITS.pdf

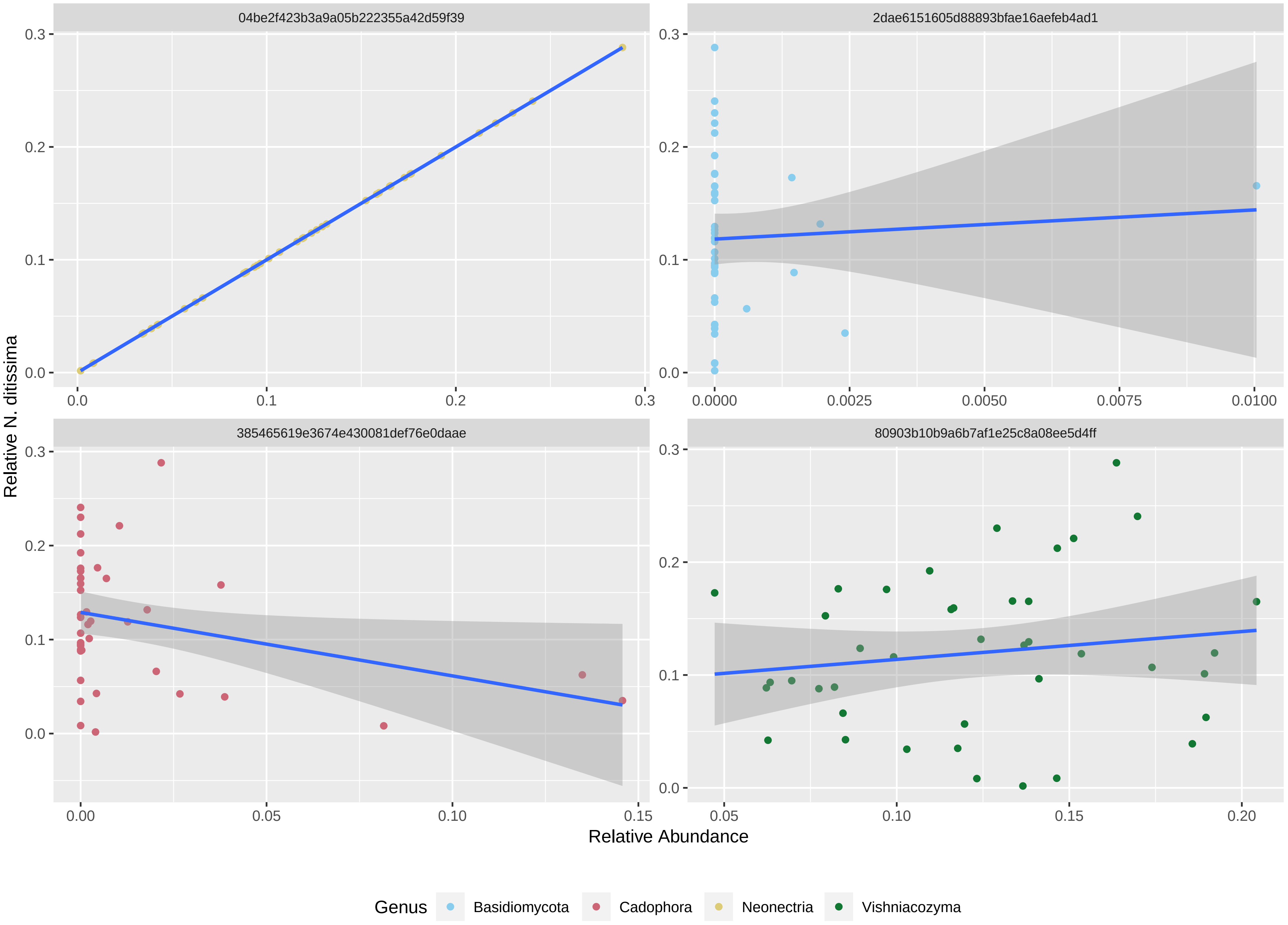

### Supplementary_Fig4A_abs_16S.pdf

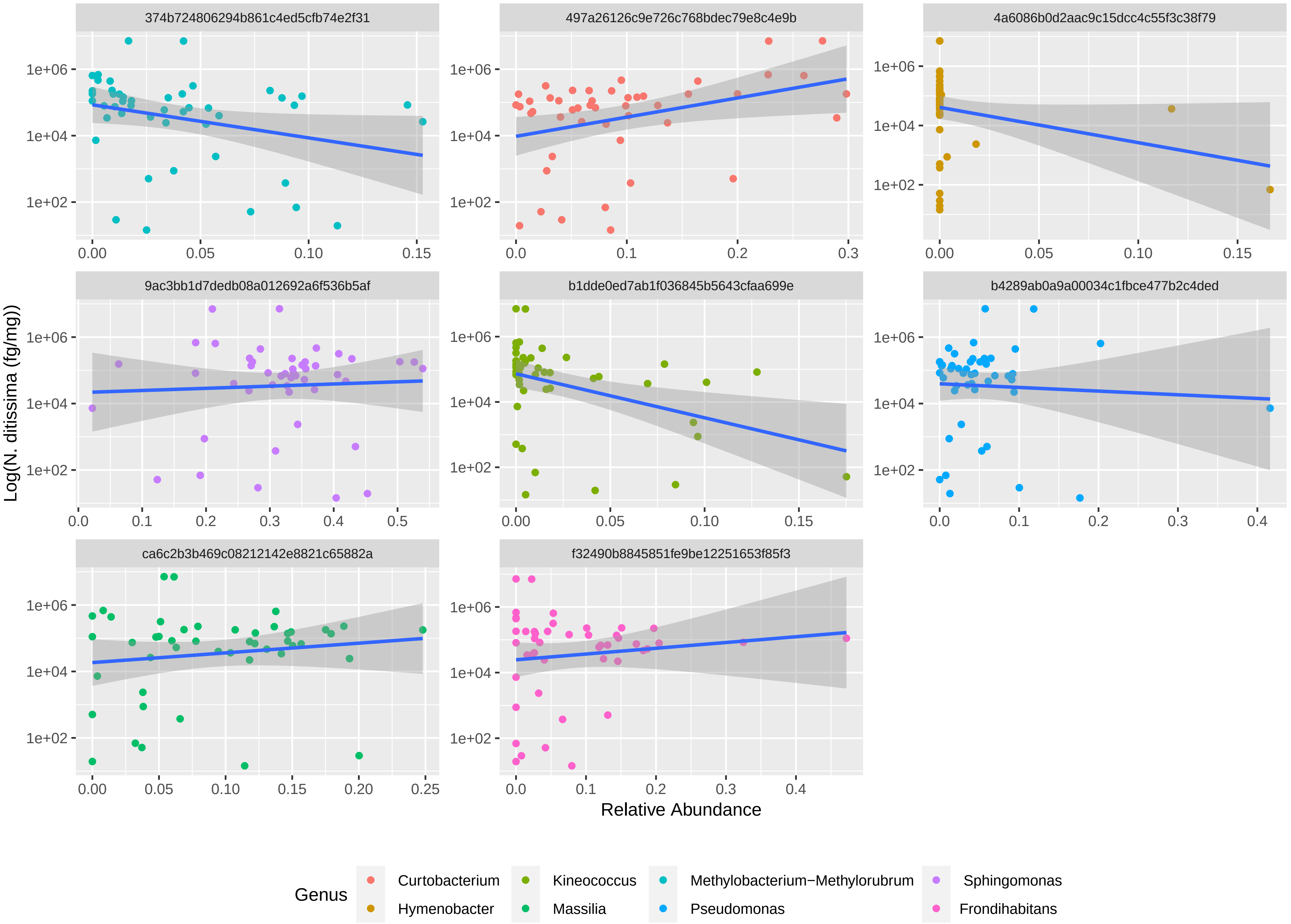

### Supplementary_Fig4B_abs_ITS.pdf

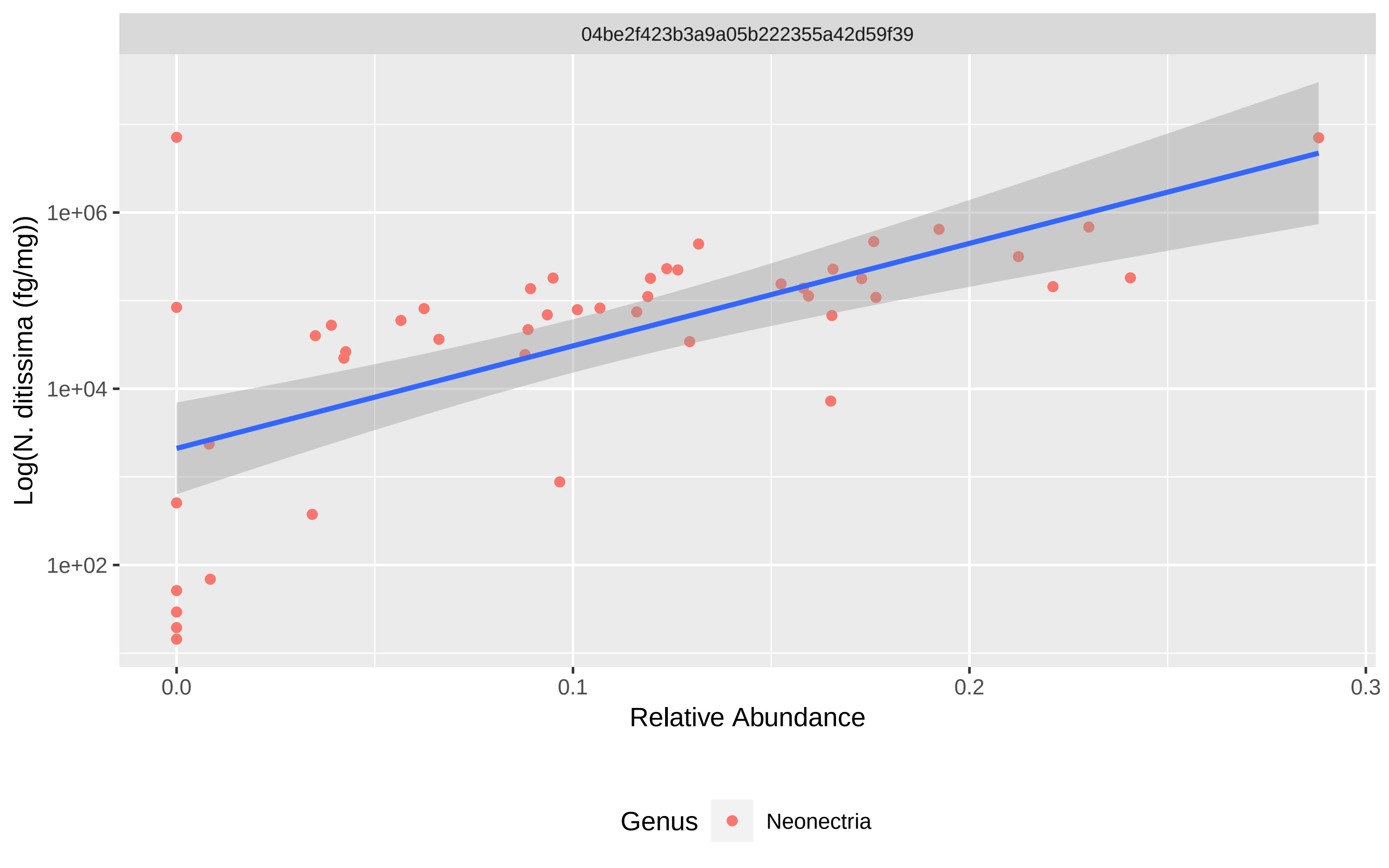
